# The Linker Domain of SNAP25 Acts as a Flexible Molecular Spacer to Ensure Efficient S-Acylation

**DOI:** 10.1101/2020.01.21.914333

**Authors:** Christine Salaun, Jennifer Greaves, Nicholas C.O. Tomkinson, Luke H. Chamberlain

## Abstract

S-Acylation of the SNARE protein SNAP25 is mediated by a subset of Golgi zDHHC enzymes, in particular zDHHC17. The ankyrin repeat (ANK) domain of this enzyme interacts with a short linear motif known as the zDHHC ANK binding motif (zDABM) in SNAP25 (112-VVASQP-117), which is downstream of the S-acylated cysteine-rich domain (85-CGLCVCPC-92). In this study, we have investigated the importance of the flexible linker (amino acids 93-111; referred to as the “mini-linker” region) that separates the zDABM and S-acylated cysteines. Shortening the mini-linker had no effect of zDHHC17 interaction but blocked S-acylation. Insertion of additional flexible glycine-serine repeats had no effect on S-acylation, whereas extended and rigid alanine-proline repeats perturbed this process. Indeed, a SNAP25 mutant in which the mini-linker region was substituted with a flexible glycine-serine linker of the same length underwent efficient S-acylation. Furthermore, this mutant displayed the same intracellular localisation as wild-type SNAP25, showing that the sequence of the mini-linker is not important in this context. By using the results of previous peptide array experiments, we generated a SNAP25 mutant predicted to have a higher affinity zDABM, and this mutant showed enhanced interaction with zDHHC17 in cells. Interestingly, this mutant was S-acylated with reduced efficiency, implying that a lower affinity interaction of the SNAP25 zDABM with zDHHC17 is optimal for S-acylation efficiency. Overall, the results of this study show that amino acids 93-111 in SNAP25 act as a flexible molecular spacer to ensure efficient coupling of enzyme-substrate interaction and S-acylation.

---

S-acylation (aka palmitoylation), the reversible attachment of fatty acids onto cysteine residues, occurs on a wide range of eukaryotic proteins [1]. This post-translational modification regulates membrane interactions of soluble proteins and mediates stabilisation and trafficking of both soluble and transmembrane proteins [2]. The occurrence of S-acylation is widespread [3, 4], with synaptic proteins apparently over-represented in the S-acylated proteome: 41 % of synaptic proteins were suggested to be modified by S-acyl chains [5]. Many of the S-acylated synaptic proteins that have been characterised are reversibly modified and undergo S-acylation cycling mediated by zDHHC S-acyltransferase and APT/ABHD thioesterase enzymes localised at synaptic regions [5–8]. However initial S-acylation of newly-synthesised synaptically-targeted proteins is likely to occur in the cell body, mediated by zDHHC enzymes localised to endoplasmic reticulum (ER) or Golgi compartments [8].

There are 23 zDHHC enzyme isoforms in the human genome [6]. The encoded enzymes are transmembrane proteins that are predominantly associated with ER and Golgi compartments [9]. The catalytic domain of zDHHC enzymes is a 51-amino acid cysteine-rich domain containing an aspartate-histidine-histidine-cysteine (DHHC) motif. The S-acylation reaction occurs via a two-step process whereby the cysteine of the DHHC motif undergoes autoacylation prior to transfer of the acyl chain to substrate cysteine [10, 11]. The catalytic cysteine of zDHHC enzymes is positioned at the cytosolic face of the membrane [12, 13] and therefore substrate cysteines must be at a similar position to allow S-acylation. Although palmitic acid is the main fatty acid used in S-acylation reactions, zDHHC enzymes can also use shorter and longer chain fatty acids as substrates [14]. The fatty acid selectivity of these enzymes is determined by specific amino acids in the transmembrane domains that determine the length of acyl chain that can be accommodated within the tepee-like cavity formed by the transmembrane helices [10, 13, 15].

Although zDHHC enzymes were recognised as S-acyltransferases over fifteen years ago [6], there is very little information available about the substrate networks of individual enzymes and how enzyme-substrate specificity is encoded. Some zDHHC enzymes, such as zDHHC3 and zDHHC7 appear to exhibit very loose substrate specificity and may have minimal interactions with their substrates [16].

Instead these enzymes may rely on a very high intrinsic activity to mediate S-acylation of any accessible cysteines at the membrane interface [16]. In contrast, some other zDHHC enzymes exhibit a much more restricted substrate network. zDHHC17 and zDHHC13 are unique in the zDHHC family as they contain an N-terminal ankyrin-repeat (ANK) domain. ANK domains are present in many different proteins and are recognised as protein interaction modules. The ANK domain of zDHHC17 is essential for S-acylation of substrates such as huntingtin, synaptosomal-associated protein of 25 kDa (SNAP25) and cysteine-string protein (CSP) [16, 17]. Our previous work identified a consensus recognition short linear motif (SLiM) in these and other substrates that mediate binding to the ANK domain of both zDHHC17 and zDHHC13 [18]. The 6-amino acid [VIAP][VIT]XXQP consensus motif makes key contacts with asparagine-100 and tryptophan-130 of zDHHC17 [19]. We named the consensus SLiM the “zDHHC Ankyrin repeat Binding Motif (zDABM)” [20]. The affinity of the SNAP25 zDABM for the ANK domain of zDHHC17 is ∼11 μM, although full-length SNAP25 has a higher affinity (0.5 μM) [19], likely owing to optimisation of binding/presentation of the zDABM to zDHHC17. The presence of other zDHHC17 binding sites in SNAP25 is unlikely as mutation of proline-117 in the zDABM blocks binding to the ANK domain, S-acylation and membrane targeting [18, 21, 22].

Using peptide arrays, we defined the sequence rules of the zDABM of SNAP25 and CSP and used these rules to predict and validate a number of new zDHHC17 interactors [20]. This analysis suggested that different zDABMs have different affinities for the ANK domain of zDHHC17 and that this can also be influenced by the identity of non-conserved residues (i.e. ‘X’ position in the zDABM) or surrounding amino acids [20]. Furthermore, using the generated sequence rules of interaction, it was possible to create zDABM sequences that displayed increased interactions with the ANK domain of zDHHC17 [20]. An interesting observation that came from this analysis is that not all of proteins that contain zDABMs are S-acylated by zDHHC17 [20], suggesting that other features of the binding partner must dictate whether or not it is an effective S-acylation substrate. Although the crystal structure of the ANK domain of zDHHC17 (with and without substrate peptide) has been reported [19], there is currently no structural information available for the full-length enzyme. Thus, it is unclear how the substrate-binding ANK domain and the catalytic DHHC domain of zDHHC17 are positioned relative to each other. Unlike other zDHHC enzymes, zDHHC17 and zDHHC13 are predicted to contain six (rather than four) transmembrane domains [23] and it is likely that the additional two transmembrane helices play some role in mediating the correct positioning of the ANK domain relative to the bilayer and DHHC domain.

Our hypothesis is that substrates of zDHHC17 require both a zDABM and also accessible cysteine residues that are appropriately positioned to allow simultaneous engagement of the zDABM with the ANK domain and the cysteines with the DHHC domain. Thus, the relative orientations and spatial separation of the ANK domain and catalytic DHHC domain of zDHHC17 must be mirrored in the zDABM and cysteines of substrate proteins to allow effective S-acylation. We also propose that the affinity of the zDABM-ANK domain interaction is suitably optimised to allow both efficient substrate engagement *and* release following S-acylation. In this study, we have tested these ideas by modifying the mini-linker region of SNAP25 that connects the zDABM and S-acylated cysteines. The results suggest that the sequence of the linker is not important for efficient S-acylation by zDHHC17 but that linker length and flexibility are important features for S-acylation. In addition, we have used knowledge about the zDABM-ANK domain interaction to generate a SNAP25 mutant which displays enhanced binding to zDHHC17. This mutant displays reduced S-acylation by zDHHC17, demonstrating the importance of binding affinity for S-acylation efficiency.

## RESULTS

### Shortening the mini-linker region between the S-acylation domain and the zDABM of SNAP25 has no effect on zDHHC17 interaction but blocks S-acylation

The linker domain of SNAP25 is the region of the protein that separates the two SNARE domains [24]. This ∼60 amino acid region includes the S-acylated cysteines (amino acids 85-92) and the zDABM (amino acids 112-117). In this study, we were interested in the region of SNAP25 separating the cysteine-rich domain and zDABM (i.e. residues 93-111) and refer to this region herein as the “mini linker” (Figure 1A).

**Figure 1.**
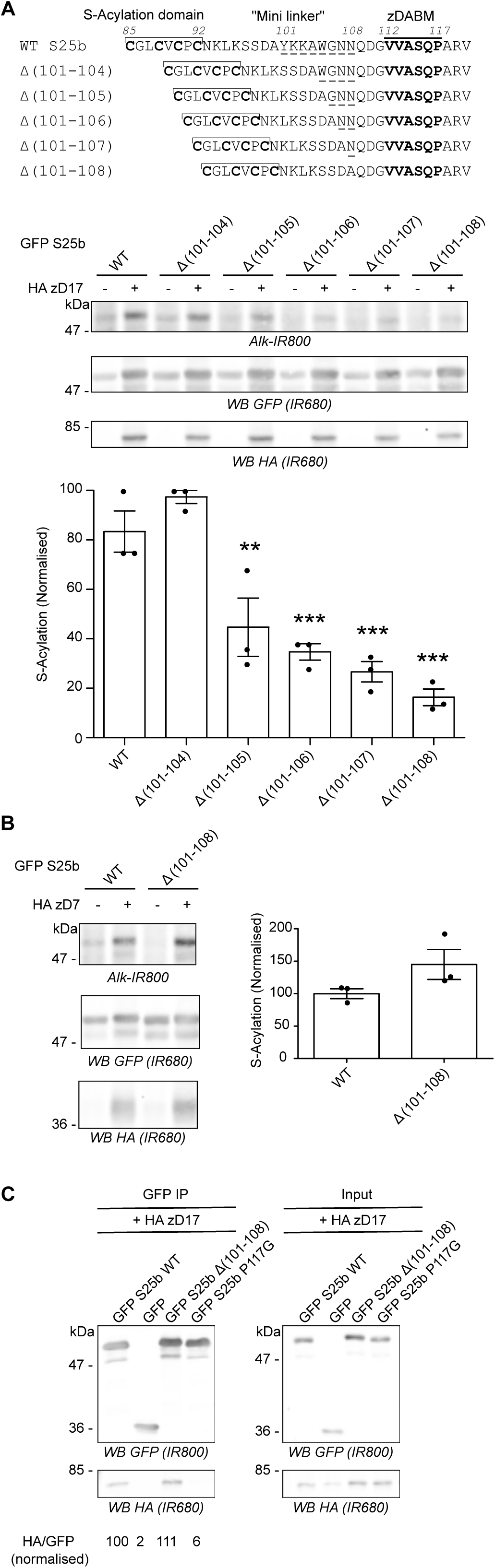
Shortening the linker region of SNAP25 decreases S-acylation by zDHHC17 without impairing interaction with the enzyme. **A.** *Top panel*: schematic of the deletions (amino acids 101 to 104, 105, 106, 107 or 108; *underlined*) introduced in the linker region of EGFP-SNAP25. *Middle and Bottom Panels*. HEK293T cells were transfected with either EGFP SNAP25b WT (GFP S25b WT) or the various deletion mutants (Δ (101-104), Δ (101-105), Δ (101-106), Δ (101-107), Δ (101-108)) together with either a plasmid encoding HA zDHHC17 (+ HA zD17) or a control pEF-BOS HA plasmid (-HA zD17). Cells were then incubated with C16:0-azide and proteins incorporating the azide fatty acid were labelled by click chemistry using an alkyne-infrared 800 dye (Alk-IR800). Isolated proteins were resolved by SDS-PAGE and transferred to nitrocellulose membranes. Representative images are shown (*Middle panel*): click chemistry signal (*Top, Alk IR800*), GFP (*Middle, IR680)* and HA (*Bottom, IR680*) immunoblots. Graph (*Bottom panel*) shows mean +/− SEM of normalised S-acylation with HA zDHHC17; *filled circles* represent individual samples (n=3 for each condition). Statistical analysis (ANOVA) showed no significant difference between the zDHHC17 mediated S-acylation of WT vs Δ (101-104) whereas there was a significant difference for WT vs Δ (101-105) (**, p<0.01), Δ (101-106), Δ (101-107) and Δ (101-108) (***, p<0.001). **B.** HEK293T cells were transfected with either EGFP SNAP25b WT (GFP S25b WT) or the Δ (101-108) deletion mutant together with HA zDHHC7 (+ HA zD7) or a control pEF-BOS HA plasmid (-HA zD7). Cells were then treated as in A. Representative images are shown on the *left panel*: click chemistry signal (*Top, IR 800*), GFP (*Middle, IR 680*), and HA (*Bottom, IR680*) immunoblots. Graph on the *right panel* shows mean +/− SEM of normalised S-acylation with HA zDHHC7; *filled circles* represent individual samples (n=3 for each condition). Statistical analysis (Student’s t-test) showed no significant difference between the S-acylation of WT vs Δ (101-108) by zDHHC7. **C.** HEK293T cells were transfected with either EGFP SNAP25b WT (GFP S25b WT), the Δ (101-108) deletion mutant (GFP S25b Δ (101-108)), or additional control plasmids (GFP SNAP25b P117G or GFP) together with HA zDHHC17 (+ HA zD17). Lysates (*right panel*, Input) were immunoprecipitated (IP) with an anti GFP antibody (*left panel*), resolved by SDS-PAGE and transferred to nitrocellulose membranes. Membranes were probed with anti GFP (*Top*) or anti HA (*Bottom*) antibodies. The ratio between the HA signal and the GFP signal was quantified for the immunoprecipitated samples and is indicated at the bottom. Position of molecular weight markers are shown on the left side of all immunoblots.

In previous work, we reported that shortening the mini-linker of SNAP25 led to a loss of membrane association induced by zDHHC17 co-expression [22]. To investigate this further, we examined S-acylation of a range of SNAP25 mutants with deletions of between 4 and 8 amino acids from the mini-linker. S-acylation was assessed by incubating transfected cells in C16:0-azide, and proteins incorporating the azide were subsequently labelled using click chemistry with an alkyne infrared dye. As shown in Figure 1A, there was a marked and significant loss of S-acylation when 5 or more amino acids were removed, with deletion of 8 amino acids having the greatest effect. This loss of S-acylation was specific to zDHHC17 and was not observed with the high activity/low selectivity enzyme, zDHHC7 (Figure 1B). These results are consistent with our previous demonstration of a loss of membrane binding of these mutants when co-expressed with zDHHC17 [22]. To examine if loss of S-acylation was associated with loss of binding to zDHHC17, we undertook immunoprecipitation (IP) experiments (Figure 1C). As shown, IP of EGFP-SNAP25 led to co-immunoprecipitation (co-IP) of HA-zDHHC17. A similar result was seen with the SNAP25 Δ(101-108) mutant, suggesting that deletion of these residues from the mini-linker has no effect on interaction of SNAP25 with zDHHC17 in HEK293T cells. In contrast, substitution of proline-117 in the zDABM of SNAP25 with a glycine residue led to a complete loss of co-IP of HA-zDHHC17, which demonstrates that this approach faithfully reports on interactions of zDHHC17 with the zDABM of SNAP25 (proline-117 is essential for this interaction) [18].

### mPEG-Click reveals the number of S-acylated cysteines on SNAP25 proteins

SNAP25 has 4 potential S-acylated cysteines. The reduction in click chemistry signal using alkyne infrared dye (Figure 1) could therefore reflect any of the following: (i) a reduction in the total number of SNAP25 molecules that are S-acylated; (ii) a reduction in the number of cysteines that are modified on individual SNAP25 molecules; or (iii) a combination of (i) and (ii). We therefore sought to implement a different click chemistry labelling approach that would allow these possibilities to be discriminated. For this, instead of reacting C16:0-azide with an alkyne infrared dye, we instead reacted it with a 5kDa PEG Alkyne (Alk-mPEG) and characterised this methodology using cells transfected with EGFP-SNAP25 and HA-zDHHC7. As can be seen in Figure 2A, this approach allowed visualisation of four distinct PEGylated bands for SNAP25 (labelled 1-4). To confirm that these immunoreactive bands represent SNAP25 modified by 1-4 mPEG groups (and hence SNAP25 S-acylated on 1-4 cysteines) we showed that: (i) these PEGylated bands were lost when all four cysteines in SNAP25 were mutated to leucines (Figure 2B); (ii) PEGylated bands were not observed with a catalytically-dead zDHHC7(C160A) mutant (Figure 2C); and (iii) there was a corresponding loss of the upper PEGylated band when one cysteine was mutated and a loss of the upper two PEGylated bands when two cysteines were mutated in SNAP25 (Figure 2D). As well as providing a methodology to more thoroughly investigate S-acylation of SNAP25, to our knowledge, this analysis also provides the first *direct* evidence that SNAP25 can be modified simultaneously on all four of its cysteines. Finally, we compared the S-acylation profile of SNAP25 when co-expressed with DHHC7 *versus* DHHC17 (Figure 2F). The WT SNAP25 protein appeared to be modified differently by zDHHC7 and zDHHC17; the amount of SNAP25 modified with 2 or 3 mPEG groups was indeed lower with zDHHC17 than with zDHHC7. mPEG-click therefore reveals that SNAP25 is less heavily S-acylated by zDHHC17 than by zDHHC7, which is consistent with zDHHC17 being a low activity / high selectivity enzyme and zDHHC7 a high activity / low selectivity enzyme [16]. Although a higher molecular weight SNAP25 band was present with zDHHC17 co-expression that migrated at the same size as SNAP25 modified by four mPEG groups, this band was present in both the absence and presence of C16:0-azide labelling (compare “- “ and “+” Az-C16:0 samples), and thus is not a PEGylated form of the protein. Instead, we believe that this is an aggregated form of SNAP25 that arises due to incomplete S-acylation by zDHHC17.

**Figure 2.**
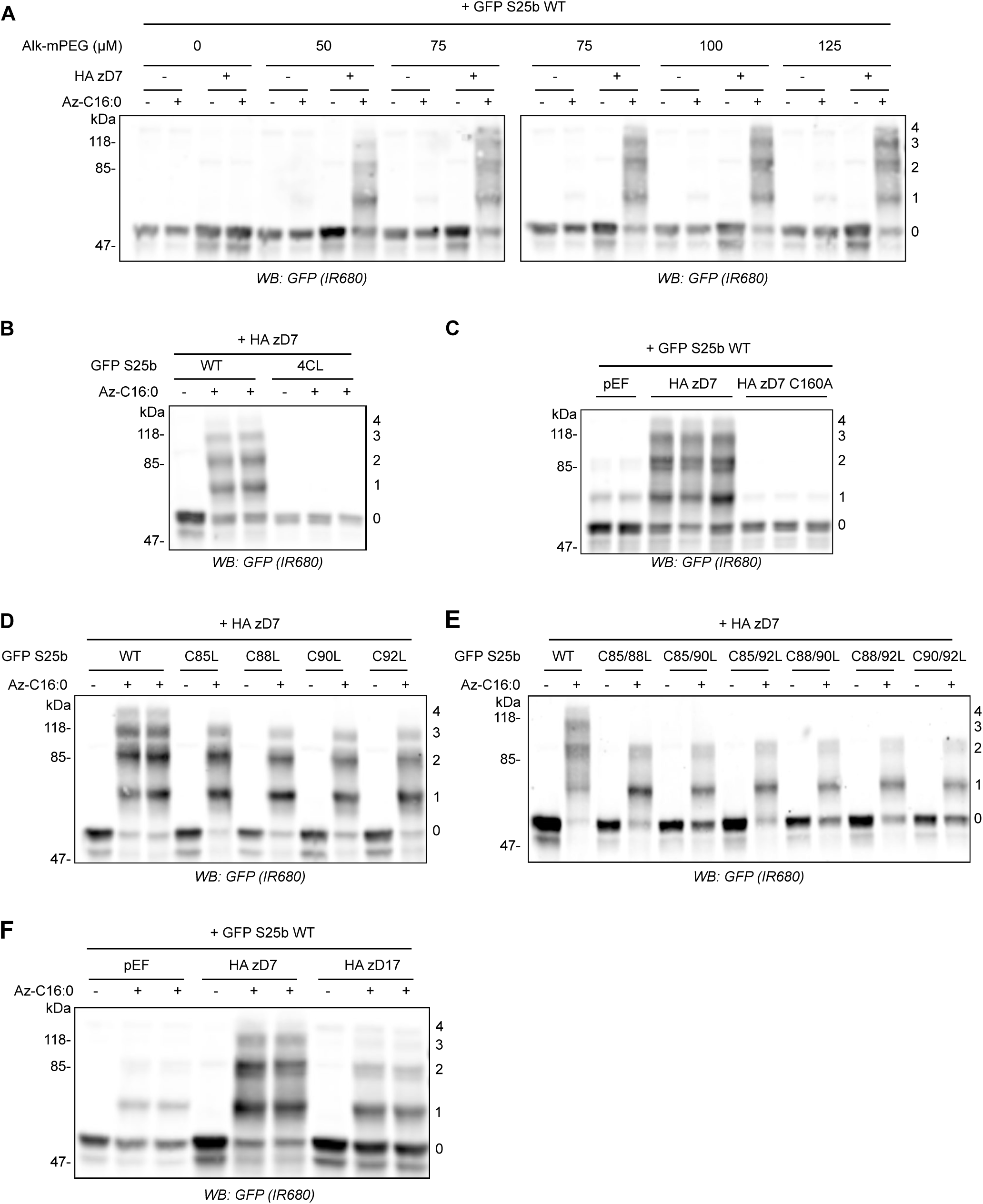
The mPEG-click technique enables the detection of the number of S-acylated cysteines of SNAP25. **A.** HEK293T cells were transfected with EGFP SNAP25b WT (GFP S25b WT) together with a plasmid encoding HA zDHHC7 (+ HA zD7) or a control pEF-BOS HA plasmid (-HA zD7). Cells were then incubated with C16:0-azide (+ Az C16:0) or palmitic acid (-Az C16:0). Proteins incorporating C16:0-azide were labelled by click chemistry using increasing concentrations of alkyne-mPEG 5K (0 to 125 μM). Isolated proteins were resolved by SDS-PAGE, transferred to nitrocellulose membranes and probed with a GFP antibody. Molecular marker weights are shown on the left whereas numbers on the right relate to the number of modified cysteines detected. **B.** HEK293T cells were transfected with either EGFP SNAP25b WT (WT) or an EGFP SNAP25b mutant where all cysteines had been mutated to leucines (4CL) together with a plasmid encoding HA zDHHC7 (+ HA zD7). Cells were then incubated with C16:0-azide (+ Az C16:0) or palmitic acid (-Az C16:0). Proteins incorporating C16:0-azide were labelled by click chemistry using 100 μM alkyne-mPEG 5K. Isolated proteins were resolved by SDS-PAGE, transferred to nitrocellulose membranes and probed with a GFP antibody. Molecular marker weights are shown on the left whereas numbers on the right relate to the number of modified cysteines detected. **C.** HEK293T cells were transfected with EGFP SNAP25b WT (GFP S25b WT) together with a plasmid encoding either HA zDHHC7 (+ HA zD7), or an inactive mutant of zDHHC7 where the catalytic cysteine had been mutated into an alanine (HA zD7 C160A) or control pEF-BOS HA plasmid (pEF). Cells were then incubated with C16:0-azide for 4 hours. Proteins incorporating C16:0-azide were labelled by click chemistry using 100 μM alkyne-mPEG 5K. Isolated proteins were resolved by SDS-PAGE, transferred to nitrocellulose membranes and probed with a GFP antibody. Molecular marker weights are shown on the left whereas numbers on the right relate to the number of modified cysteines detected. **D.** HEK293T cells were transfected with EGFP SNAP25b WT (WT) or single cysteine mutants (C85L, C88L, C90L and C92L) together with a plasmid encoding HA zDHHC7 (+ HA zD7). Cells were then labelled and analysed as in panel *B*. **E.** HEK293T cells were transfected with EGFP SNAP25b WT or double cysteine mutants (C85/88L, C85/90L, C85/92L, C88/90L C88/92L and C90/92L) together with a plasmid encoding HA zDHHC7 (+ HA zD7). Cells were then labelled and analysed as in panel *B*. **F.** HEK293T cells were transfected with EGFP SNAP25b WT together with a plasmid encoding HA zDHHC7 (HA zD7), HA zDHHC17 (HA zD17) or a control plasmid pEF-BOS HA (pEF). Cells were then labelled and analysed as in panel *B*.

We then studied the S-acylation profile of EGFP-SNAP25 WT and mini-linker deletion mutants when co-expressed with zDHHC7 or - 17 using the newly characterised mPEG-Click technique. The mPEG-click results were consistent with the results obtained with the alkyne infrared dye click technique, and the deletions had little effect on S-acylation catalysed by zDHHC7 (Figure 3A). In contrast, for S-acylation mediated by zDHHC17, the progressive shortening of the linker region induced an equally progressive disappearance of the PEGylated bands, consistent with a reduction in S-acylation (Figure 3A). Our hypothesis is that this loss of S-acylation occurs due to reduced accessibility of cysteines in SNAP25 to the active site of zDHHC17 (i.e. that the binding site and catalytic site of zDHHC17 are separated by a minimal distance, which also requires a minimal separation of the zDABM and cysteine-rich domain of SNAP25). We reasoned that cysteines in SNAP25 that are closer to the zDABM (e.g. C92) should become inaccessible to the zDHHC17 active site before cysteines that are further away from the zDABM (e.g. C85) as the mini-linker is shortened. To test this, we introduced C-to-L mutations within two of the deletion mutants that showed a partial but not complete loss of S-acylation by zDHHC17 (Δ(101-105) and Δ(101-106); Figure 1A). Either the first cysteine of the S-acylation domain (C85) or the last one (C92) were mutated to leucine. Interestingly, the C85L mutation had more impact on the S-acylation profile of the deletion mutants than the C92L mutation (Figure 3B), showing that C85 is more efficiently S-acylated within the deletion mutants than C92. This suggests that C92 becomes more inaccessible for S-acylation when the mini-linker sequence is shortened, consistent with it being closer to the zDABM zDHHC17 binding site.

**Figure 3.**
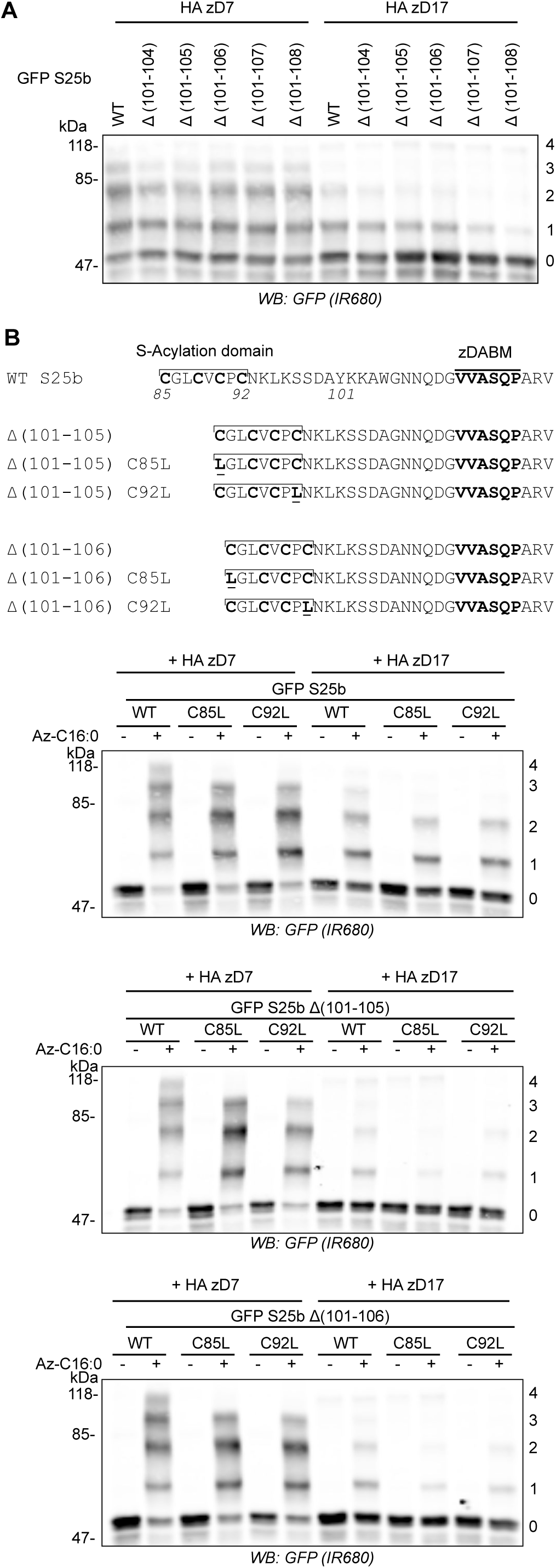
mPEG-Click analysis of SNAP25 linker deletion mutants. **A.** HEK293T cells were transfected with either EGFP SNAP25b WT (GFP S25b WT) or the various deletion mutants (Δ (101-104), Δ (101-105), Δ (101-106), Δ (101-107), Δ (101-108)) together with a plasmid encoding either HA zDHHC7 (HA zD7) or HA zDHHC17 (HA zD17). Cells were then incubated with C16:0 azide and proteins incorporating the fatty acid azide were labelled by click chemistry using an alkyne-mPEG 5K. Isolated proteins were resolved by SDS-PAGE and transferred to nitrocellulose membranes and probed with a GFP antibody. Position of molecular weight markers are shown on the left whereas numbers on the right relate to the number of modified cysteines in SNAP25. **B.** *Top Panel.* Schematic of the mutants of EGFP SNAP25b used in the analysis. Cysteine-to-Leucine mutations (underlined) were introduced at position 85 (C85L) or 92(C92L) of EGFP SNAP25b WT (WT S25b) or the deletion mutants EGFP SNAP25b Δ (101-105) and Δ (101-106). *Lower panels.* HEK293T cells were transfected with the various constructs described above together with a plasmid encoding either HA zDHHC7 (HA zD17) or HA zDHHC17 (HA zD17). Cells were then incubated with C16:0 azide (+ Az C16:0) or palmitic acid (-Az C16:0). Proteins incorporating the fatty acid azide were labelled by click chemistry using alkyne-mPEG 5K. Isolated proteins were resolved by SDS-PAGE, transferred to nitrocellulose membranes and probed with a GFP antibody. Position of molecular weight markers are shown on the left of all immunoblots whereas numbers on the right relate to the number of mPEG-modified cysteines.

### Efficient S-acylation of SNAP25 is disrupted by introducing structure into the mini-linker sequence

As S-acylation was disrupted by introducing short deletions to the mini-linker region of SNAP25, we investigated if inserting additional amino acids was also disruptive. The zDABM and mini-linker region of SNAP25 are disordered [18, 25] and when disorder was maintained through the insertion of a 10-amino acid (G4S)_2_ flexible sequence (GGGGSGGGGS) [26], there was no effect on S-acylation mediated by zDHHC7 and a very slight but significant increase in S-acylation by zDHHC17 (Figure 4A). In contrast, when a more rigid amino acid sequence was introduced at the same position within SNAP25, consisting of Ala-Pro repeats (SNAP25b (AP)_2_, (AP)_4_ and (AP)_6_) [26], there was a clear effect on S-acylation mediated by zDHHC17 but not zDHHC7 (Fig 4B). Specifically, the insertion of the 8 amino acid sequence (AP)_4_ or of the 12 amino acid sequence (AP)_6_ significantly decreased SNAP25 S-acylation, consistent with a displacement of the S-acylation site from the active site of zDHHC17.

**Figure 4.**
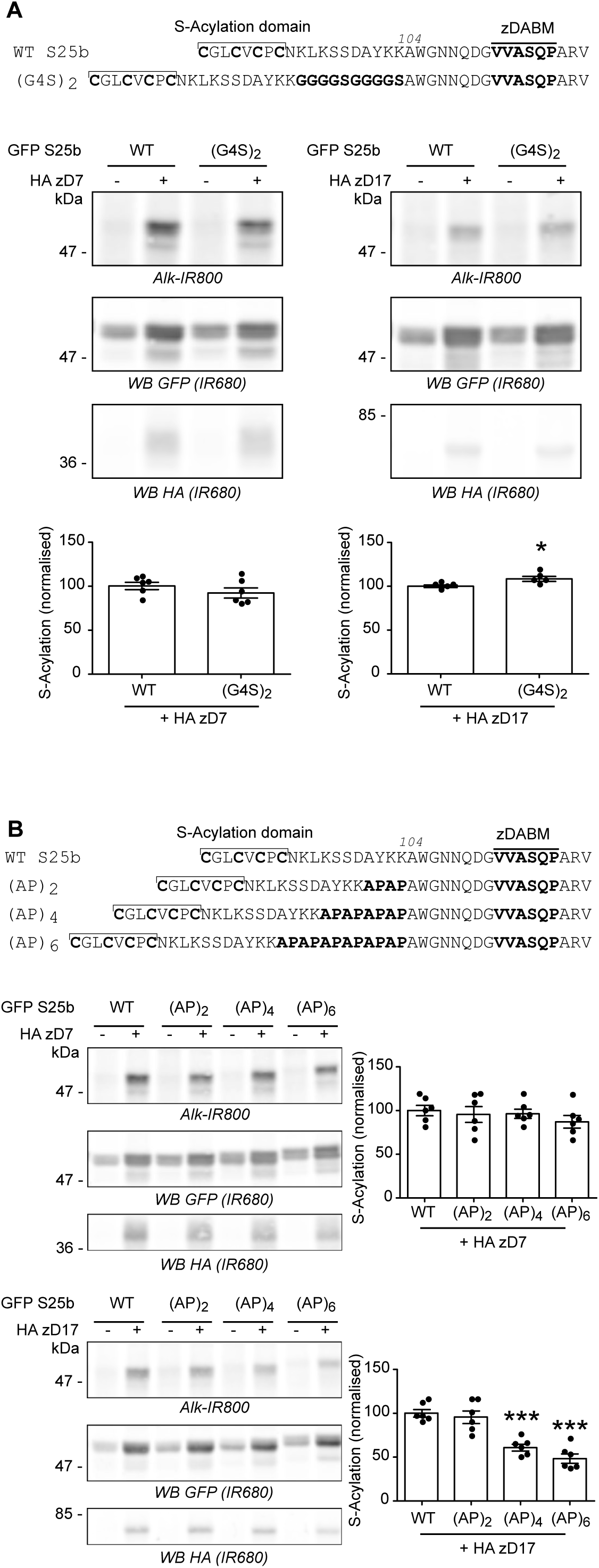
Differential effects of flexible *versus* rigid sequences inserted into the linker domain of SNAP25. **A.** *Top panel*. Schematic showing the introduction of a duplicated flexible sequence composed of 4 glycines and 1 serine (G4S)_2_ at position 104 of EGFP SNAP25b. *Middle and bottom panels.* HEK293T cells were transfected with either EGFP SNAP25b WT (GFP S25b WT) or the (G4S)_2_ insertion mutant together with a plasmid encoding either HA zDHHC7 (+ HA zD7), HA zDHHC17 (+ HA zD17) or a control pEF-BOS HA plasmid (-HA zD7, - HA zD17). Cells were then incubated with C16:0 azide and proteins incorporating the fatty acid azide were labelled by click chemistry using an alkyne-800 infrared dye (Alk-IR800). Isolated proteins were resolved by SDS-PAGE and transferred to nitrocellulose membranes. Representative images are shown (*Middle panel)*: click chemistry signal (*Top, Alk IR800*), GFP (*Middle, IR680*) and HA (*Bottom, IR680*) immunoblots. Graphs (*Bottom panel*) show mean +/− SEM of normalised S-acylation with HA zDHHC7 or HA zDHHC17; *filled circles* represent individual samples (n=6 for each condition). Statistical analysis (Student’s t-test) showed no significant difference between the S-acylation of WT vs (G4S)_2_ by zDHHC7 whereas there was a significant difference of the S-acylation of WT vs (G4S)_2_ by zDHHC17 (*, p<0.05). **B.** *Top panel*. Schematic showing the introduction of (AP)_2_, (AP)_4_ or (AP)_6_ at position 104 of EGFP SNAP25b. *Middle and bottom panels.* HEK293T cells were transfected with either EGFP SNAP25b WT (GFP S25b WT) or the (AP) insertion mutants ((AP)_2_, (AP)_4_ and (AP)_6_) together with a plasmid encoding HA zDHHC7 (+ HA zD7), HA zDHHC17 (+ HA zD17) or a control pEF-BOS HA plasmid (-HA zD7, - HA zD17). Cells were then incubated with C16:0-azide and proteins incorporating the fatty acid azide were labelled by click chemistry using an alkyne-800 infrared dye. Isolated proteins were resolved by SDS-PAGE and transferred to nitrocellulose membranes. Representative images are shown (*Left panel)*: click chemistry signal (*Top, IR800*), GFP (*Middle, IR680*) and HA (*Bottom, IR680*) immunoblots. Graphs (*Right panel*) show mean +/− SEM of normalised S-acylation with HA zDHHC7 or HA zDHHC17; *filled circles* represent individual samples (n=6 for each condition). Statistical analysis (ANOVA) showed no significant difference between the S-acylation of WT *versus* any of the (AP) insertion mutants by zDHHC7 whereas there was a significant difference of the S-acylation of WT *versus* (AP)_4_ and (AP)_6_ by zDHHC17 (***, p<0.001). Position of molecular weight markers are shown on the left side of all immunoblots.

Together with the previous data showing that shortening of the mini-linker region is detrimental for zDHHC17-mediated S-acylation of SNAP25, this suggest that the distance between the S-acylation domain and the zDABM sequence of SNAP25b is critical for its S-acylation by zDHHC17, and that whereas additional flexible amino acid sequences are tolerated, rigid sequences that introduce structural constraints are inhibitory.

### Replacing the mini-linker region with a flexible linker does not affect SNAP25 S-acylation by zDHHC17

The previous analyses have shown that the length of the mini-linker sequence is an important factor for efficient S-acylation of SNAP25 by zDHHC17. To examine if the sequence of the mini-linker is also important, this region (93-111) was entirely replaced with a glycine-serine (GS) linker of the same length (Figure 5A). Unexpectedly, this mutant showed a partial loss of S-acylation mediated by zDHHC7, which might reflect a difference in membrane affinity (see Discussion). However, this replacement linker sequence had no effect on SNAP25 S-acylation by zDHHC17 (Figure 5B). Accordingly, we found that the GS mutant could co-IP zDHHC17 as efficiently as WT SNAP25, showing that the interaction between SNAP25 and zDHHC17 is not dependent on the exact sequence of the mini-linker region (Figure 5C).

**Figure 5.**
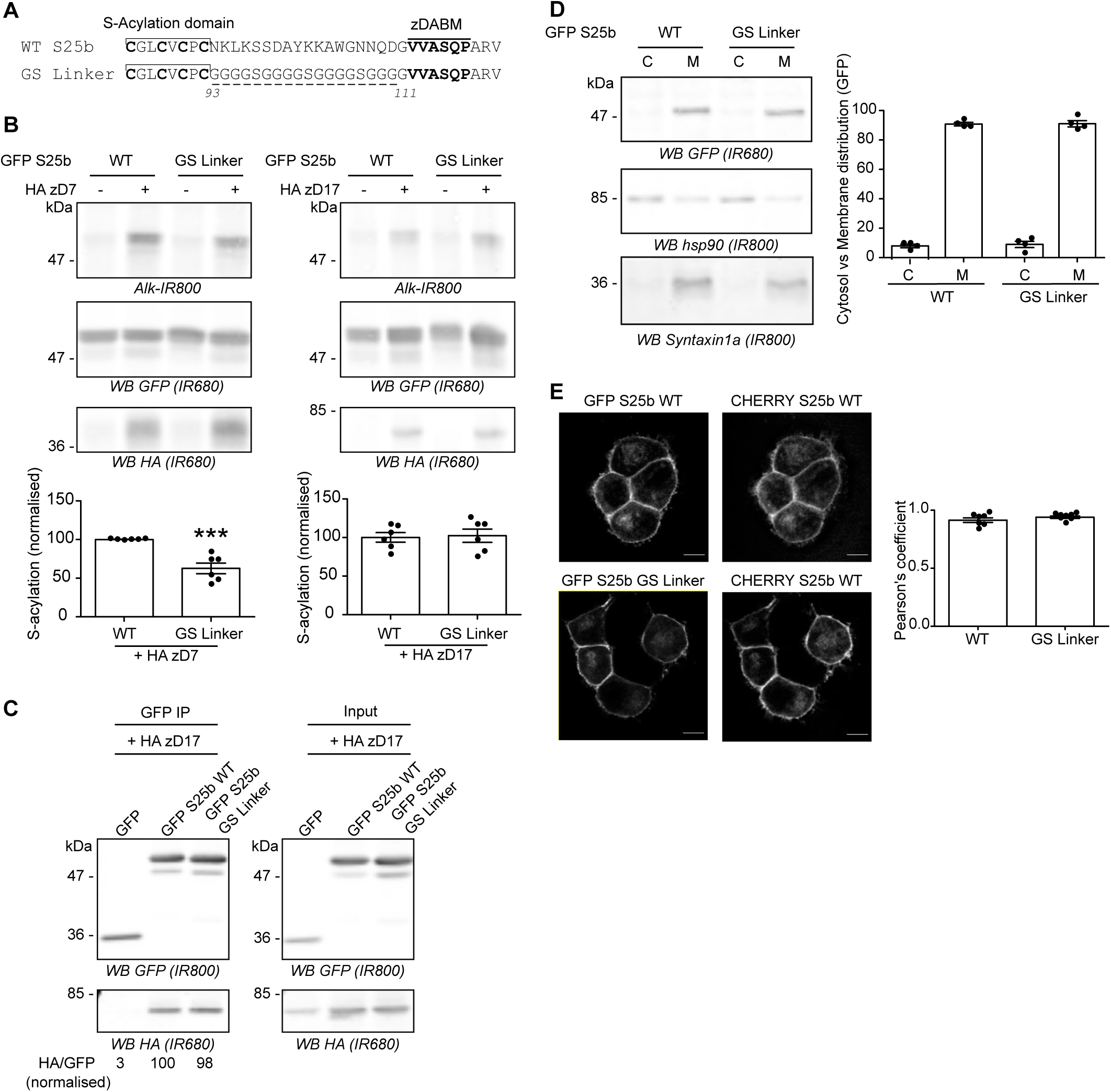
Replacing the linker region of SNAP25 with a flexible glycine-serine linker of the same length has no effect on SNAP25 S-acylation by zDHHC17 in HEK293T cells or its localisation in PC12 cells. **A.** Amino acids between the S-acylation domain and the zDABM of EGFP SNAP25b (WT S25b) were replaced by several copies of the flexible GGGGS sequence to create the GS Linker mutant. **B.** HEK293T cells were transfected with either EGFP SNAP25b WT (GFP S25b WT) or the GS Linker mutant together with either a plasmid encoding either HA zDHHC7 (+ HA zD7), HA zDHHC17 (+ HA zD17) or a control pEF-BOS HA plasmid (-HA zD7, - HA zD17). Cells were then incubated with C16:0-azide and proteins incorporating the fatty acid azide were labelled by click chemistry using an alkyne-800 infrared dye. Isolated proteins were resolved by SDS-PAGE and transferred to nitrocellulose membranes. Representative images are shown: click chemistry signal (*Top, Alk IR800*), GFP (*Middle, IR680*), and HA (*Bottom, IR680*) immunoblots. Graphs (*bottom panel*) show mean +/− SEM of normalised S-acylation with HA zDHHC7 (HA zD7) or HA zDHHC17 (HA zD17); *filled circles* represent individual samples (n=6 for each condition). Statistical analysis (Student’s t-test) showed a significant difference between the S-acylation of WT *versus* the GS Linker mutant by zDHHC7 (***, p<0.001) whereas there was no significant difference of the S-acylation of WT *versus* the GS Linker mutant by zDHHC17. **C.** HEK293T cells were transfected with either EGFP SNAP25b WT (GFP S25b WT), GS Linker mutant (GFP S25 GS Linker) or a GFP control plasmid together with HA zDHHC17 (+ HA zD17). Lysates (*Right panel*, Input) were immunoprecipitated (IP) with a GFP antibody (*Left panel*), separated by SDS-PAGE and transferred to nitrocellulose membranes. Membranes were probed with GFP (*Top, IR800*) and HA (*Bottom, IR680*) antibodies. The ratio between the HA signal and the GFP signal was quantified for the immunoprecipitated samples and is indicated at the bottom. **D.** PC12 cells were transfected with either EGFP SNAP25b WT (WT) or the GS Linker mutant (GS Linker). Cells were fractioned into cytosol fraction (C) and membrane (M) fractions. Samples were separated by SDS-PAGE and transferred to nitrocellulose membranes. Representative images are shown (*Left panel*): GFP (*Top, IR680*), Hsp90 (*Middle, IR800*) and Syntaxin1a (*Bottom, IR800*). The graph (*Right panel*) shows mean +/− SEM of the cytosol *versus* membrane distribution of the EGFP-tagged proteins; *filled circles* represent individual samples (n=4 for each condition). Statistical analysis (Student’s t-test) showed no significant difference in the distribution of both constructs. **E.** PC12 cells were transfected with either EGFP SNAP25b WT (GFP S25 WT) or the GS Linker mutant (GFP S25 GS Linker) together with a mCHERRY SNAP25b WT (CHERRY S25 WT). The *middle* panel shows the localisation of the WT mCHERRY-tagged SNAP25 and the *left* panel the expression of the co-expressed WT or mutant EGFP-tagged SNAP25 proteins. Scale bars represent 5 μm. The graph (*Right panel*) represents mean +/− SEM of the Pearson’s co-localisation coefficient of the GFP signal *versus* mCHERRY signal for both conditions; *filled circles* represent individual images (n=7 for each condition). Statistical analysis (Student’s t-test) showed no significant difference between the two constructs. Position of molecular weight markers are shown on the left side of all immunoblots.

Although the replacement linker had no effect on S-acylation by zDHHC17, it is possible that it could impact the localisation of SNAP25. To examine the effect on localisation, we performed experiments in PC12 cells. EGFP-SNAP25 is efficiently S-acylated by endogenous zDHHC enzymes in this cell line, and mutation of the zDABM blocks S-acylation, demonstrating dependence on endogenous zDHHC17 [22, 27]. In addition, SNAP25 is endogenously-expressed in PC12 cells and over-expressed EGFP-SNAP25 localises correctly to the plasma membrane and endosome/*trans* Golgi network compartment in these cells [28]. Cellular fractionation showed that the GS Linker mutant associated with membranes to the same extent as wild-type SNAP25 in PC12 cells (Figure 5D). Moreover, confocal imaging and analysis also revealed that the mini-linker mutations had no effect on the intracellular localisation: both EGFP-tagged SNAP25 WT and GS Linker mutant co-localised to the same extent with a mcherry-tagged SNAP25 protein when co-expressed in PC12 cells (Figure 5E).

Altogether these results show that the exact amino acid sequence of the mini-linker region of SNAP25 is not important for its S-acylation by zDHHC17 or subsequent targeting to the plasma membrane and endosomal compartments.

### Increasing the affinity of the zDABM for zDHHC17 leads to reduced S-acylation efficiency

Lemonidis *et al* [20] have previously defined the potentially optimal zDABM amino acid sequence for the binding of substrates to the zDHHC17 ANK domain. A ‘favourable peptide’ sequence was synthesized based on the sequence rules defined for the SNAP25 zDABM and found to interact more strongly than a WT SNAP25 zDABM peptide with the soluble ANK domain of zDHHC17 [20]. We therefore modified the zDABM of SNAP25 to include most of the amino acids of the ‘favourable peptide’ (mutant referred to herein as ‘LFP’ for Lemonidis Favourable Peptide) (Figure 6A). We found that these amino acid substitutions indeed led to a marked increase in co-IP of HA-zDHHC17 (Figure 6B), confirming that the strength of interaction was increased. When S-acylation of this high affinity mutant was examined, it was found that there was no effect on S-acylation mediated by zDHHC7, whereas the LFP substitutions significantly decreased the S-acylation of SNAP25 by zDHHC17 (Figure 6C), showing that a higher affinity interaction between the enzyme and substrate was actually detrimental for S-acylation of SNAP25 in cells.

**Figure 6.**
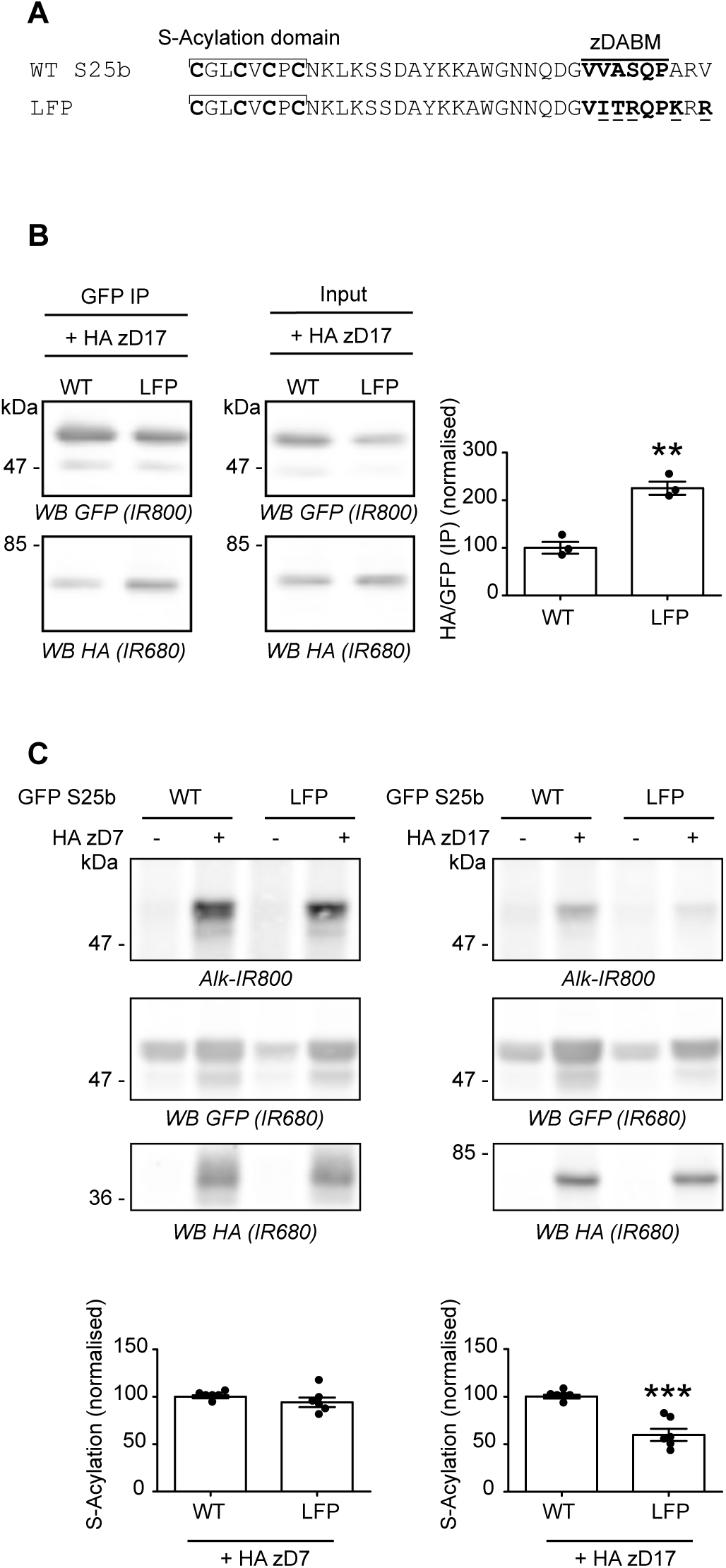
Increasing the affinity of SNAP25b for zDHHC17 results in decreased S-acylation. **A.** Amino acids within the zDABM of SNAP25 were mutated (underlined) according to the Lemonidis Favourable Peptide (LFP) sequence to increase its affinity for zDHHC17. **B.** HEK293T cells were transfected with either EGFP SNAP25b WT (WT) or the LFP mutant (LFP) together with HA zDHHC17 (+ HA zD17). Lysates (*Input*) were immunoprecipitated (IP) with a GFP antibody, separated by SDS-PAGE and transferred to nitrocellulose membranes. Representative images are shown on the *left panel.* Membranes were probed with GFP (*Top, IR800*) or HA (*Bottom, IR680*) antibodies. The ratio between the HA signal and the GFP signal was quantified for the immunoprecipitated samples. The graph (*Right panel*) shows mean +/− SEM of this ratio; *filled circles* represent individual samples (n=3 for each condition). Statistical analysis (Student’s t-test) showed a significant difference between the interaction of zDHHC17 with SNAP25 WT *versus* the LFP mutant (**, p<0.01). **C.** HEK293T cells were transfected with either EGFP SNAP25b WT (GFP S25b WT) or the LFP mutant (LFP) together with either a plasmid encoding HA zDHHC7 (+ HA zD7), HA zDHHC17 (+ HA zD17) or a control pEF-BOS HA plasmid (-HA zD7, - HA zD17). Cells were then incubated with C16:0-azide and proteins incorporating the fatty acid azide were labelled by click chemistry using an alkyne-800 infrared dye. Isolated proteins were resolved by SDS-PAGE and transferred to nitrocellulose membranes. Representative images are shown (*Top panel*): click chemistry signal (*Top, Alk IR800*), GFP (*Middle, IR680*) and HA (*Bottom, IR680*) immunoblots. Graphs (*bottom panel*) show mean +/− SEM of normalised S-acylation with HA zDHHC7 (+ HA zD7) or HA zDHHC17 (+ HA zD17); *filled circles* represent individual samples (n=6 for each condition). Statistical analysis (Student’s t-test) showed no significant difference between the S-acylation of WT *versus* LFP by zDHHC7 whereas there was a significant difference of the S-acylation of WT *versus* LFP by zDHHC17 (***, p<0.001). Position of molecular weight markers are shown on the left side of all immunoblots.

## DISCUSSION

Identification of the substrate networks of individual zDHHC enzymes is a major requirement for progress of the S-acylation field. Experiments to determine these substrate networks (e.g. by zDHHC depletion and quantitative proteomics) will be complicated by the observation that zDHHC enzymes can have overlapping substrate selectivity [1]. SNAP25, the focus of the current study, interacts with and is S-acylated by zDHHC17 [6, 16, 18, 20, 22, 27]. However, co-expression experiments have revealed that this protein can also be modified by zDHHC3, zDHHC7 and zDHHC15 [6, 21]. Indeed, zDHHC3 and zDHHC7 S-acylate SNAP25 to a greater extent than zDHHC17, despite zDHHC3/7 displaying no detectable interaction with SNAP25 [16]. Furthermore, zDHHC3/7 can modify a diverse set of substrates that have no obvious sequence or structural similarities. Based on this, we believe that zDHHC3/7 are “high activity/low selectivity” enzymes that modify accessible and reactive cysteines on proteins that co-localise at the Golgi (e.g. during trafficking of newly-synthesised proteins), in the absence of any specific enzyme-substrate recognition. In contrast, zDHHC17 is a “low activity/high selectivity” enzyme that depends upon specific interactions with its substrate proteins to mediate S-acylation. This idea is supported by several observations, including: (i) removal of the substrate-binding ANK domain prevents S-acylation by zDHHC17 [16] and (ii) mutation of the zDABM motif of SNAP25 prevents S-acylation by zDHHC17 [18, 20–22]. Indeed, it is interesting to note that although zDHHC3 and zDHHC7 robustly S-acylate SNAP25 in co-expression experiments, mutation of the zDABM in SNAP25 leads to a loss of membrane binding (and thus presumably S-acylation) in PC12 cells [21], demonstrating the importance of zDHHC17 for SNAP25 S-acylation in this cell type at least.

The current study sought to determine features of SNAP25 that are important for the coupling of binding to zDHHC17 with subsequent S-acylation. This question is important as we showed that zDHHC17 interacts with zDABM sequences from a diverse set of proteins and that several of these binders are not known to be S-acylated [20]. Thus, we were interested in the features of SNAP25 that allow it to be a zDHHC17 S-acylation substrate. The results of the study showed that there is a minimum separation length required between the cysteine-rich domain and zDABM for S-acylation by zDHHC17. These two regions of SNAP25 are separated by 19 amino acids (residues 93-111); as this part of SNAP25 is thought to be disordered [25], an extended peptide sequence of this length would be in the region of 7.6 nm (assuming 0.4 nm per amino acid [29]). Removal of four amino acids was tolerated, whereas removal of five or more amino acids from this linker region reduced S-acylation, suggesting a minimal required separation in the order of 6 nm between zDABM and cysteines for efficient S-acylation. Although the structure of full-length zDHHC17 has not been reported, these approximate calculations of the displacement of the ANK domain and DHHC domain of zDHHC17 can be tested when structural information is available. By using mPEG-click analysis of C85L and C92L mutants of Δ(101-105) and Δ(101-106), it was found that the C85L mutation had a greater effect than the C92L mutation in the context of the deletion mutants, whereas there was no obvious difference between these mutations in full-length SNAP25. This supports that idea that as the mini-linker is shortened, cysteines closer to the zDABM (i.e. C92) become less accessible for S-acylation before cysteines further from the zDABM (i.e. C85), thus why C85 mutation has a greater effect than C92 mutation on S-acylation of the linker deletion mutants.

The mPEG-click technique newly described here was used to generate additional data to the ones obtained with the widely used dye-click technique. Indeed mPEG-click reveals the number of S-acylated cysteines and also differentiates between mutations affecting the overall S-acylation of a protein and mutations affecting the S-acylation of specific cysteines within a protein. Coupled with pulse chase and time course analyses, mPEG-click could be used to study the dynamics and processivity of cysteine S-acylation / deacylation within multiply S-acylated proteins. Since the S-acylation profile of a protein is simply revealed with antibodies against this protein, overexpression of the protein of interest is not strictly necessary as long as the antibody is sensitive enough to detect the endogenous protein, and that the epitopes are not masked by the addition of the mPEG group to the S-acylated cysteines. In our opinion, mPEG-click might provide an easy and straightforward mean to investigate the S-acylation status of any endogenous or overexpressed protein in various cell types.

In contrast to mutations that shortened the mini-linker region, addition of a 10-amino acid flexible glycine-serine linker had no major effect on S-acylation. This suggests that whereas there is a strict requirement for a minimal separation distance between zDABM and S-acylated cysteines, there is more flexibility around the maximal length, presumably because a longer disordered and flexible linker can adopt a spatial orientation that can simultaneously dock the zDABM and cysteines of a substrate into the ANK domain and DHHC domain of zDHHC17, respectively. In contrast, the insertion of rigid inflexible alanine-proline sequences into the mini-linker region of SNAP25 perturbed S-acylation by zDHHC17 presumably by limiting spatial flexibility of the linker region and hence the spatial optimisation of zDABM and cysteine-rich domain of this protein.

Interestingly, analysis of the GS Linker mutant that had a complete replacement of the 19-amino acid linker with a similar length GS linker suggested that the mini-linker sequence is not important for either S-acylation or intracellular targeting of SNAP25. It was notable however that this GS Linker mutant was less efficiently S-acylated by zDHHC7. Our previous work suggested that the hydrophobic cysteine-rich domain of SNAP25 (residues 85-92) plays an important role in initial membrane interaction of SNAP25 with Golgi membranes prior to S-acylation [22]. In contrast, a study by the Lang group suggested a role for positively-charged amino acids in the (large) linker domain in mediating this initial membrane interaction [30]. However, it is worthwhile further reflecting on the proposed role of positively-charged amino acids in initial membrane interaction of SNAP25 [30]. The results of the current study clearly show that lysine residues in the mini-linker are dispensable for S-acylation mediated by zDHHC17 and for membrane association and intracellular targeting in PC12 cells. Furthermore, any positively-charged residues upstream of the cysteine-rich domain are also dispensable as the minimal S-acylation and membrane targeting domain in SNAP25b is amino acids 85-120 [31]. Thus, it is likely that any role played by positively-charged amino acids of SNAP25 is secondary to the prominent role of the hydrophobic cysteine-rich domain in initial membrane targeting of this protein [22]. Indeed, the importance of the non-acylated cysteine-rich domain in membrane interactions of SNAP25 was further shown by a recent study [32]. The effect of mini-linker replacement on S-acylation by zDHHC7 may reflect the idea that this enzyme does not interact with SNAP25 directly [16] but rather mediates “stochastic” S-acylation of this protein when it is present at the membrane. Thus, a slight loss in membrane affinity caused by mini-linker replacement could lead to the reduced S-acylation seen with zDHHC7, whereas a slight reduction in membrane affinity may still be sufficient to promote efficient zDHHC17-SNAP25 interaction and subsequent S-acylation.

The affinity of the zDABM of SNAP25 for the ANK domain of zDHHC17 was calculated to be ∼ 11 μM, although full-length SNAP25 had a higher affinity [19]. This interaction affinity is clearly sufficient to allow robust isolation of the protein complex by co-IP. However we were interested in whether enhancing the affinity of interaction would lead to higher levels of S-acylation. To test this idea, we used data from a previous study by our group [20] to generate a SNAP25 mutant, which was predicted to have an increased affinity for the ANK domain of zDHHC17. In support of this idea, the mutant captured higher levels of zDHHC17 by co-IP. Despite this increased binding capacity, the LFP high affinity mutant displayed a reduced S-acylation by zDHHC17, suggesting that tight binding between zDHHC enzymes and other proteins is not always conducive to efficient S-acylation; perhaps tighter binding limits the catalytic turnover of the enzyme.

This study demonstrates that the length and flexible character of the mini-linker region of SNAP25 are important features for efficient S-acylation. In contrast, the sequence of the mini-linker is less important for efficient S-acylation as long as it confers structural flexibility. Future work will determine if the sequence of the mini-linker is important for the function of SNAP25 in fusion pore dynamics and exocytosis [33].

## EXPERIMENTAL PROCEDURES

### Antibodies

Mouse GFP antibody (Clontech, clone JL8, used at 1:4,000) was obtained from Takara (Saint-Germain-en-Laye, France). Rat HA antibody (Roche, clone 3F10, used at 1:1,000) was from Sigma (Poole, UK). Rabbit HSP90 antibody (C45G5, used at 1:1,000) was from Abcam (Cambridge, UK). Rabbit Syntaxin1a antibody (110 302, used at 1:1,000) was from Synaptic Systems (Gottingen, Germany). IR dye conjugated secondary antibodies (used at 1:20,000 dilution) were purchased from Li-COR Biosciences (Cambridge, UK).

### Plasmid DNA

cDNA encoding human zDHHC7 and zDHHC17, and the (AP)_6_ and GS Linker SNAP25b mutants were synthesized by Thermo Fisher Scientific (Loughborough, UK). The cDNA encoding zDHHC7 and zDHHC17 were sub-cloned into pEF-BOS HA [6], whereas the SNAP25b constructs were sub-cloned in pEGFP-C2 [22]. Wild-type rat SNAP25b (in pEGFP-C2), SNAP25 mutants with deletions of the linker region and mcherry-SNAP25 were previously described [22, 28]. All other mutants described in this report were generated by site-directed mutagenesis using oligonucleotide primers synthesised by Sigma (Poole, UK). The validity of all constructs was confirmed by sequencing (Dundee DNA sequencing Service, UK).

### Cells

HEK293T cells (CRL-3216, ATCC) were grown in DMEM media (Thermo Fisher Scientific) supplemented with 10% foetal bovine serum. PC12 cells (CRL-1721, ATCC) were grown in Advanced RPMI 1640 media (Thermo Fisher Scientific) containing 10% horse serum, 5% foetal bovine serum and 1% glutamine. All cells were grown at 37°C in a humidified atmosphere containing 5% CO_2_.

### Cell transfection

For substrate S-acylation and immunoprecipitation assays, HEK293T were plated on poly-D-lysine coated 24-well plates (Corning BioCoat, VWR, UK) and transfected with 0.33 μg of EGFP-SNAP25 plasmid and 0.66 μg of pEF-BOS HA plasmid (either empty as a control or encoding the zDHHC enzymes). 2 μl of polyethylene imine (PEI) (1 mg / ml stock) (linear PEI mw 25,000, #43896, Alpha Aesar, UK) was added to the DNA mix, incubated for 20 min and then added to the cells, which were analysed the following day.

For cell fractionation and immunofluorescence analyses, PC12 cells were plated on poly-D-lysine coated 24-well plates or on 12 mm BD poly-D-lysine coated coverslips (Thermo Fisher Scientific, UK), respectively. Cells were transfected using Lipofectamine 2000 (Invitrogen, UK) with a ratio of 2 μl of Lipofectamine / μg of DNA. For fractionation experiments, 1 μg of EGFP SNAP25 WT or GS Linker mutant was transfected whereas 0.2 μg of EGFP SNAP25b WT or GS Linker mutant was transfected together with 0.2 μg of mCHERRY SNAP25b WT for immunofluorescence analysis. Cells were incubated with the transfection mixture for approximately 48 h before analysis.

### Cell Labelling with fatty acids

HEK293T cells in 24 well plates were incubated with 100 μM of either palmitic acid (P0500, Sigma, UK) or C16:0-azide [15] in 350 μl of serum free DMEM supplemented with 1 mg/ml defatted BSA (A7031, Sigma, UK) for 4 h at 37°C.

### Detection of palmitate labelled probes in S-acylated proteins

Cells were washed once with PBS then lysed on ice in 100 μl of 50 mM Tris pH 8.0 containing 0.5% SDS and protease inhibitors (P8340, Sigma, UK). Conjugation of IR800CW Alkyne Dye or mPEG5K Alkyne to C16:0-azide was carried out for 1 h at room temperature with end-over-end rotation by adding an equal volume (100 μl) of freshly prepared click chemistry reaction mixture containing the following: 5 μM IRDye 800CW Alkyne (929-60002, Li-COR, UK) or 200 μM mPEG5k-Alkyne (JKA3177, Sigma, UK), 4 mM CuSO_4_ (451657, Sigma, UK), 400 μM Tris[(1-benzyl-1*H*-1,2,3-triazol-4-yl)methyl]amine (678937, Sigma, UK) and 8 mM ascorbic acid (A15613, Alpha Aesar, UK) in dH_2_O. 67 μl of 4x SDS sample buffer containing 100 mM DTT was then added to the 200 μl sample. Protein samples were incubated at 95°C for 5 min and 19 μl was resolved by SDS-PAGE and transferred to nitrocellulose for immunoblotting analysis. Immunoblots were quantified with the ImageStudio software from Li-Cor. Graphs and statistical analysis was performed using GraphPad software.

### Immunoprecipitation

Cells were co-transfected with plasmids encoding EGFP-SNAP25 constructs together with a HA-tagged zDHHC17 construct on 24 well plates. Three identical wells were gathered for each immunoprecipitation. Cells were washed, lysed in 200 μl of PBS, 0.5% Triton X-100 (T8787, Sigma, UK) supplemented with protease inhibitors for 30 min on ice and clarified by centrifugation at 16,000 g for 10 min at 4°C. The supernatants were then adjusted to 0.2% Triton X-100 by the addition of 300 μl of cold PBS. Whereas 50 μl of lysate was kept as the input fraction, 450 μl of lysate was added to 8 μl (slurry volume) of washed GFP-Trap agarose beads (GTA-20, Chromotek GmbH, Germany). The tubes were incubated on an end-over-end rotator at 4°C for 1 h. The beads were then washed twice with PBS and eluted with 50 μl of 2x SDS PAGE sample buffer containing 50 mM DTT with a 5 min incubation at 95°C.

### Fractionation of PC12 cells

Transfected PC12 cells were detached from the wells in 500 μl PBS and transferred to Eppendorf tubes. Cell fractionation was performed as described by S. Baghirova *et al* [34]. Briefly, cells were washed in PBS, resuspended in 150 μl of Buffer A (150 mM NaCl, 50 mM HEPES pH 7.4, 1M Hexylene glycol (112100, Sigma, UK), 25 μg/ml digitonin (D141, Sigma, UK)) supplemented with protease inhibitors and rotated end-over-end at 4°C for 10 min. The supernatant of a 2,000 g, 10 min centrifugation at 4°C was collected as the ‘Cytosolic Fraction’. The pellet was washed briefly in Buffer A before being resuspended by vortexing in 150 μl of Buffer B (150 mM NaCl, 50 mM pH 7.4, 1M Hexylene glycol, 1% v/v Igepal (CA-630, Sigma, UK) supplemented with protease inhibitors. After 30 min incubation on ice, the lysates were centrifuged at 7,000 g for 10 min at 4°C; the supernatant was collected as the ‘Membrane Fraction’. Both Cytosolic and Membrane fractions were supplemented with 4x SDS PAGE loading buffer containing 100 mM DTT and incubated at 95°C for 5 min. Proteins were resolved by loading an equal volume of each fraction by SDS-PAGE and analysis by immunoblotting. The effectiveness of the membrane/cytosol fractionation was assessed by probing with antibodies against Syntaxin1a and Hsp90.

### Immunofluorescence, confocal microscopy and image analysis

Transfected cells were washed once with PBS and fixed in 4% formaldehyde for 30 min. The cells were then washed again in PBS and mounted on glass slides in Mowiol. All microscopy analysis was performed on a Leica SP8 confocal microscope; image stacks were acquired with the Lightning function. A single slice was chosen as a representative image. Image quantification was performed with the Fiji software. The Co-localisation Threshold tool was used to calculate the Pearson’s co-localisation coefficient *r*. Statistical analysis was performed with the GraphPad software.

## ACKNOWLEDGEMENTS

We are grateful to Graeme MacKenzie (SIPBS imaging facility) for advice with confocal microscopy. This work was funded by grants from the BBSRC (BB/L022087/1) and the MRC (MR/R011842/1).

